# Recombination and lineage-specific mutations linked to the emergence of SARS-CoV-2

**DOI:** 10.1101/2020.02.10.942748

**Authors:** Juan Ángel Patiño-Galindo, Ioan Filip, Ratul Chowdhury, Costas D. Maranas, Peter K. Sorger, Mohammed AlQuraishi, Raul Rabadan

## Abstract

The emergence of SARS-CoV-2 underscores the need to better understand the evolutionary processes that drive the emergence and adaptation of zoonotic viruses in humans. In the betacoronavirus genus, which also includes SARS-CoV and MERS-CoV, recombination frequently encompasses the Receptor Binding Domain (RBD) of the Spike protein, which, in turn, is responsible for viral binding to host cell receptors. Here, we find evidence of a recombination event in the RBD involving ancestral linages to both SARS-CoV and SARS-CoV-2. Although we cannot specify the recombinant nor the parental strains, likely due to the ancestry of the event and potential undersampling, our statistical analyses in the space of phylogenetic trees support such an ancestral recombination. Consequently, SARS-CoV and SARS-CoV-2 share an RBD sequence that includes two insertions (positions 432-436 and 460-472), as well as the variants 427N and 436Y. Both 427N and 436Y belong to a helix that interacts directly with the human ACE2 (hACE2) receptor. Reconstruction of ancestral states, combined with protein-binding affinity analyses using the physics-based trRosetta algorithm, reveal that the recombination event involving ancestral strains of SARS-CoV and SARS-CoV-2 led to an increased affinity for hACE2 binding, and that alleles 427N and 436Y significantly enhanced affinity as well. Structural modeling indicates that ancestors of SARS-CoV-2 may have acquired the ability to infect humans decades ago. The binding affinity with the human receptor was subsequently boosted in SARS-CoV and SARS-CoV-2 through further mutations in RBD. In sum, we report an ancestral recombination event affecting the RBD of both SARS-CoV and SARS-CoV-2 that was associated with an increased binding affinity to hACE2.

**Importance:** This paper addresses critical questions about the origin of the SARS-CoV-2 virus: what are the evolutionary mechanisms that led to the emergence of the virus, and how can we leverage such knowledge to assess the potential of SARS-like viruses to become pandemic strains? In this work, we demonstrate common mechanisms involved in the emergence of human-infecting SARS-like viruses: first, by acquiring a common haplotype in the RBD through recombination, and further, through increased specificity to the human ACE2 receptor through lineage specific mutations. We also show that the ancestors of SARS-CoV-2 already had the potential to infect humans at least a decade ago, suggesting that SARS-like viruses currently circulating in wild animal species constitute a source of potential pandemic re-emergence.

## Introduction

Within one year since it was first reported in mid-December 2019, the COVID-19 pandemic had already caused more than 2.3 million fatalities and over 80 million non-lethal cases of severe respiratory disease worldwide^1^. The causative agent of COVID-19, SARS-CoV-2, was a previously unknown RNA coronavirus (CoV) of the betacoronavirus genus^2^, with 80% similarity at the nucleotide level to the Severe Acute Respiratory Syndrome coronavirus (SARS-CoV)^3^. SARS-CoV and SARS-CoV-2 are still the only members of *Sarbecovirus* subgenus of betacoronavirus known to infect humans. Other members of this subgenus are frequently found in bats, which are hypothesized to be the natural reservoir of many zoonotic coronaviruses^4^. In January 2020, a viral isolate from a *Rhinolophus affinis* bat obtained in 2013 from the Yunnan Province in China (named RaTG13) was reported to have 96% similarity to SARS-CoV-2^5^. Shortly thereafter, another viral strain also collected from bat (*Rhinolophus mayalanus*), RmYN02, was reported to have 97.2% similarity to SARS-CoV-2 in the ORF1ab gene but only 61.3% identity in the RBD (93.3% genome sequence identity). These discoveries suggested that the ancestors of the outbreak virus may have been circulating in a recent past in bats^6^. Additional surveillance of wild populations identified two lineages of CoVs in pangolins that were also similar to SARS-CoV-2: one obtained from animals sampled in Guangxi in 2017, the other sampled in Guangdong in 2019. Genome-wide, these pangolin viruses were more distant from SARS-CoV-2 than RaTG13 or RmYN02, having approximately 90% sequence similarity, although the Guangdong lineage was the closest relative of SARS-CoV-2 with respect to the Receptor Binding Domain (RBD) of the Spike protein. Consequently, pangolins were postulated as possible intermediate hosts^7^. However, to date, the specific molecular and evolutionary events that enable viruses such as the recent ancestors of SARS-CoV-2 to jump species remain poorly characterized.

For a virus to infect a new host species it must have the capacity to cross the host cell membrane and interact productively with the cell’s biochemical machinery. One of the primary factors determining the tropism of Sarbecoviruses is the RBD, a region in the Spike protein that binds to the host receptor; for both SARS-CoV and SARS-CoV-2 this receptor is angiotensin I converting enzyme 2 (ACE2)^8,9,10^. Other cellular factors may also contribute to the adaptation of a virus to a new host, including their ability to use cellular proteases (such as TMPRSS2^9^ in the case of SARS-CoV-2, as well as furin^11,12^) as well as their ability to avoid or exploit host immune responses.

The ability of viral populations to emerge in new hosts are influenced by factors such as rapid mutation rates and recombination^13^ which lead to both high genetic variability and high rates of evolution (estimated in coronaviruses to be between 10^-4^ and 10^-3^ substitutions per site per year)^14^. Previous genome-wide analyses in coronaviruses have estimated that their evolutionary rates are of the same order of magnitude as in other fast-evolving RNA viruses^15,16^. Recombination in RNA viruses, which is known to be frequent in coronaviruses, can lead to the acquisition of genetic material from other viral strains^17^. Indeed, recombination has been proposed to have played a major role in the generation of new coronavirus lineages such as SARS-CoV^17^. In particular, prior studies have suggested that SARS-CoV-2 may have undergone recombination with other members of the *Sarbecovirus* subgenus^2^, possibly including its closest relatives, RaTG13 and pangolin-CoVs^18^, but other studies have yielded contradictory results^19^. Experimental analyses have revealed that the RNA proofreading exoribonuclease (nps14-ExoN) is a key mediator of recombination in CoVs^20^.

In this work, we reconstruct the evolutionary events that have accompanied the emergence of SARS-CoV-2, with a special emphasis on the RBD and its adaptation for binding to its receptor, human ACE2 (hACE2). In particular, we identify a specific recombination event which involved ancestral lineages to both SARS-CoV and SARS-CoV-2. By reconstruction of ancestral states of the Spike gene, we find that this ancestral recombination event is associated with an increased binding affinity towards hACE2. Through structure-based binding affinity predictions, we infer that affinity further increased in the lineages leading to SARS-CoV and SARS-CoV-2. These observations suggest that RBD recombination provides the backbone for CoV adaptation to humans, with subsequent point mutations further permitting high affinity RBD-hACE2 binding.

## Results

### Recombination hotspots in betacoronavirus

To understand how recombination contributes to the evolution of betacoronaviruses across different viral subgenera and hosts, we analyzed 45 betacoronavirus sequences from the five major subgenera that infect mammals (*Embevovirus, Merbecovirus, Nobecovirus, Hibecovirus* and *Sarbecovirus*) (Supplementary Table 1a)^21^. We identified 103 recombination events by using the RPDv4 package^22^, with additional validation provided by ensuring that recombination events were phylogenetically informative and that they had a different evolutionary history than the rest of the genome, (Figure 1a, Methods). Enrichment analysis showed that recombination often involves the N-terminus of the Spike protein, which includes the RBD (adjusted *p*-val. < 10^-4^, binomial test on sliding window of 800 nucleotides) (Figure 1b, Supplementary Figure 1a). Enrichment for recombination events persists even after we restricted the analysis to the most common host (bats), suggesting that recombination was not driven simply by sampling of multiple human sequences (Supplementary Figure 1b). We conclude that recombination in betacoronaviruses frequently involves the Spike protein across viral subgenera and hosts.

**Fig. 1.**
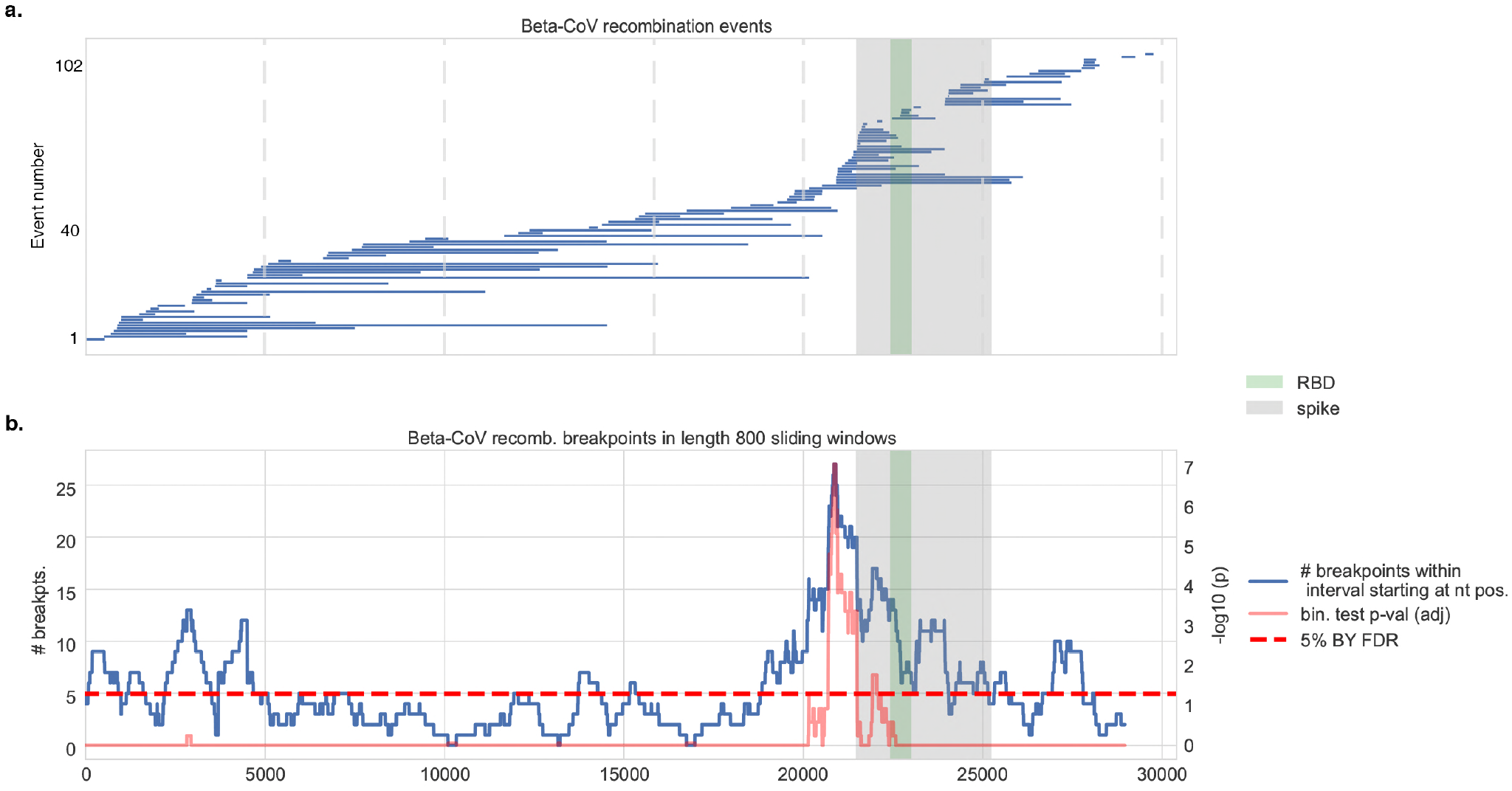
Recombination analysis of betacoronaviruses. **a.** Distribution of 103 inferred recombination events among human and non-human beta-CoV isolates showing the span of each recombinant region along the viral genome with respect to SARS-CoV coordinates. The spike protein and its RBD are highlighted. **b.** Sliding window analysis shows (blue curve) the distribution of recombination breakpoints (either start or end) in 800 nucleotide (nt) length windows upstream (namely, in the 5’ to 3’ direction) of every nt position along the viral genome. The spike protein, and in particular the RBD and its immediate downstream region, are significantly enriched in recombination breakpoints in betacoronaviruses. Benjamini-Yekutieli (BY) corrected *p*-values are shown (red curve), and the 5% BY FDR is shown for reference (dotted line).

### MERS-CoV recombination frequently involves the Spike gene

To study how recombination affects emerging human betacoronaviruses viruses at the level of individual viral species, we initially focused our attention on the Middle East respiratory syndrome coronavirus (MERS-CoV). This virus has been extensively sampled in humans and in bactrian camels (*Camelus bactrianus ferus*), which are recognized as the source of recent zoonosis^23^. Using 381 MERS-CoV sequences (170 from human, 209 from camel and 2 from bat) (Supplementary Table 1b) we found that the Spike region overlaps with the genome segment in which the majority of recombination segments took place (83% or 20 of 24 identified events) (Figure 2a) with enrichment of detectable recombination breakpoints in the Spike and Membrane genes (Figure 2b, Supplementary Figure 1c). Enrichment was not observed when the analysis was restricted to human MERS-CoV samples only (n=170) possibly due to the lower number and diversity of sequences available (Supplementary Figure 2). Thus, the enrichment of recombination events involving the Spike gene is also observed at a viral species level.

**Fig. 2.**
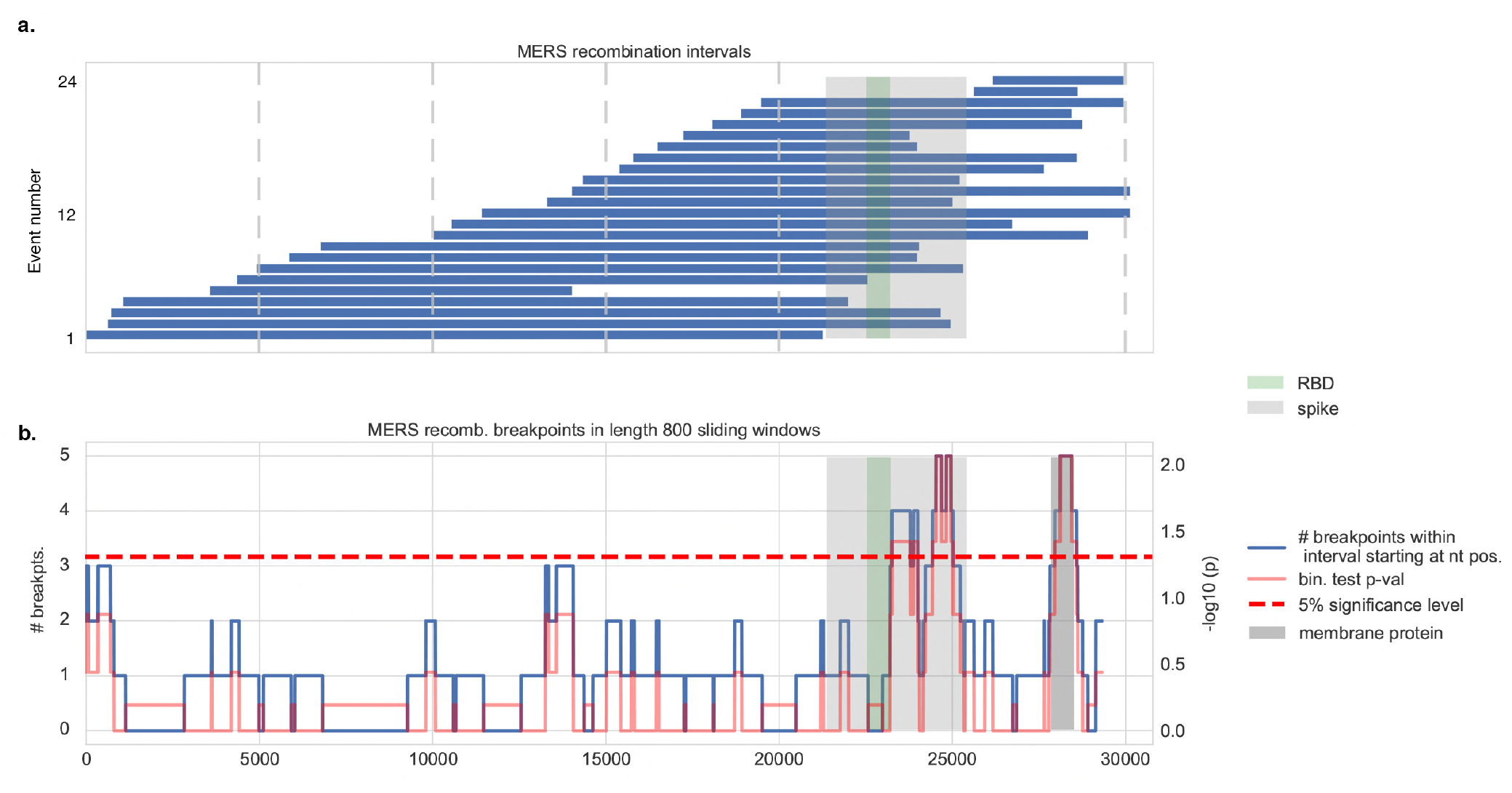
Recombination analysis in MERS coronaviruses. **a.** Distribution of 24 recombination events among human and non-human MERS-CoV isolates. The spike protein and its RBD are highlighted. **b.** Sliding window analysis shows (blue curve) the distribution of recombination breakpoints (either start or end) in 800 nucleotide (nt) length windows upstream (namely, in the 5’ to 3’ direction) of every nt position along the viral genome. The spike protein, and the RBD in particular, overlap with widows that are enriched in recombination breakpoints. Binomial test *p*-values (red curve) and the 5% significance level are shown (dotted line). The MERS-CoV membrane protein is highlighted (dark gray); it also shows an enrichment of recombination breakpoints.

### Identification of an RBD recombination event involving ancestors of SARS-CoV-2 and SARS-CoV

We then asked if any signal of recombination could be found in the recent history of SARS-CoV-2. We performed recombination analysis with the RDP4 package and detected a statistically significant recombination event in the RBD (at genome positions 22614-23032) involving ancestral lineages to both SARS-CoV-2 and SARS-CoV (*p*-val. < 0.003, with four different recombination tests – see Online Methods). This phylogenetically informative event (unresolved quartets: 13%) passed the Expected Likelihood Weight test and the Shimodaira-Hasegawa tree topology test as well (*p*-val. < 0.001), indicating that the RBD recombinant region displayed an evolutionary history significantly distinct from the evolutionary history of the rest of the genome. SARS-CoV and its closest related strains form a well-supported cluster with SARS-CoV-2, RaTG13 and the pangolin CoVs in the RBD. However, they are more distant in the rest of the genome (Figures 3a, 3b). Finally, using the space of phylogenetic trees (also known as the BHV^24^ space), we devised a randomization test that quantifies the impact of specific taxa or clades on the divergence between two different evolutionary histories (see Phylogenetic analyses). We found that the SARS-CoV/SARS-CoV-2 clade from the RBD phylogeny (Figure 3a) had the largest impact on the divergence between the whole-genome and the RBD trees of Sarbecoviruses, when compared to other clades and randomly sampled subsets of taxa (*p*-val. < 0.0001, Supplementary Figure 3). Together, these results support our hypothesis that a recombination event at RBD positions 22614-23032 involved viruses ancestral to the SARS-CoV and SARS-CoV-2 lineages.

**Fig. 3.**
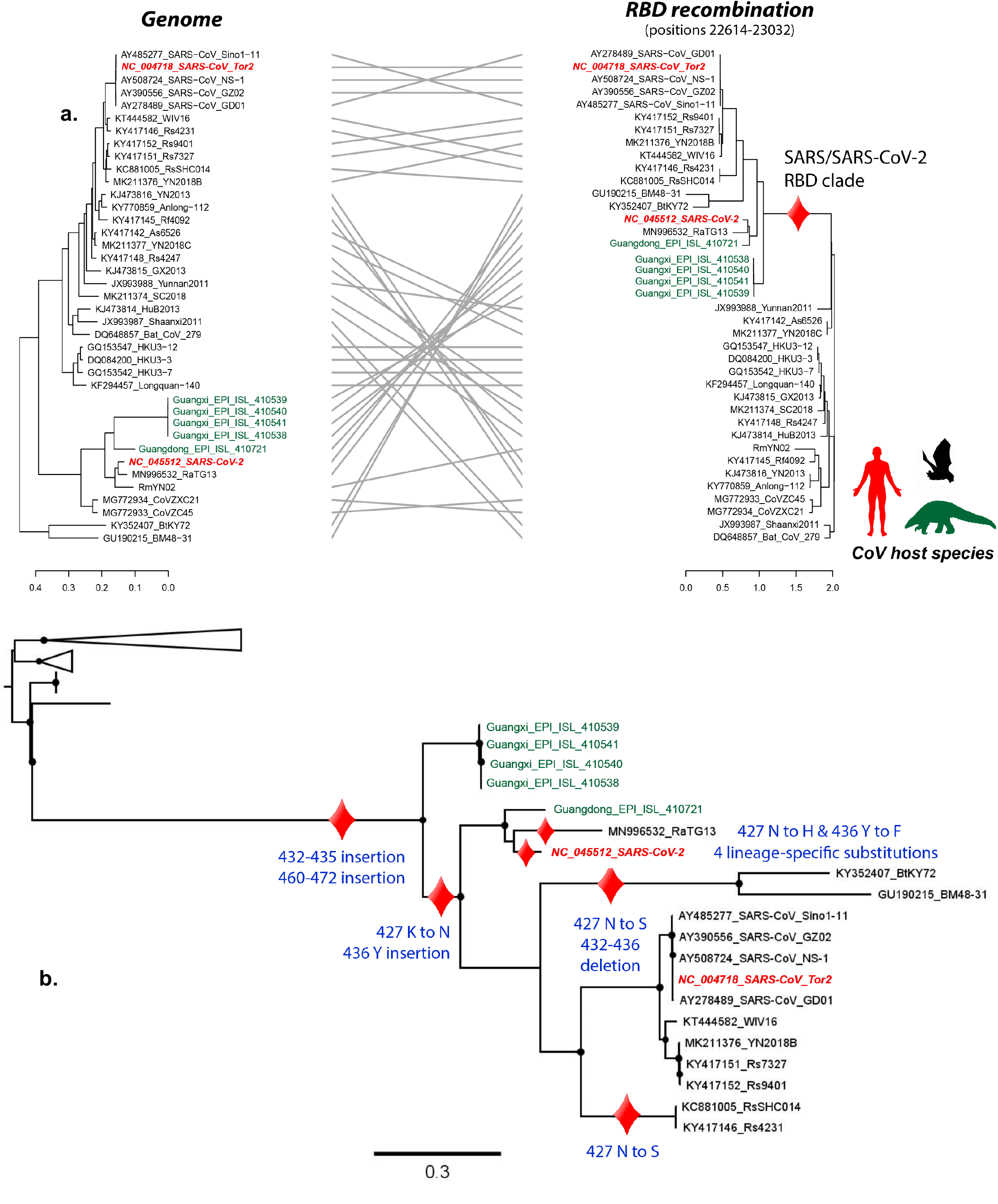
Tracing the evolution of RBD in Sarbecoviruses. Ancestral reconstruction analyses were performed using the maximum likelihood phylogenetic tree derived from RBD recombination event as input. a) Track of the evolutionary changes that occurred in the RBD from humaninfecting Sarbecoviruses and their closest relatives. Black circles in the ML tree represent nodes with Shimodaira-Hasegawa-like support higher than 0.80. b) Distribution of Likelihood values associated with the most likely amino acid inferred for the different MRCAs in the RBD ancestral reconstruction analysis. Red dots represent positions highlighted in our work (Spike 333, 359, 427, 436, 432-435, 460-472, 484, 505).

### Tracing the evolutionary changes in RBD

We traced the evolutionary changes that occurred in the SARS/SARS-CoV-2 clade (from the RBD phylogeny, Figure 3a) through an ancestral state reconstruction analysis based on a maximum-likelihood approach. Our analyses revealed that the evolution of the RBD in this clade is characterized by two insertions at spike positions 432-436 and 460-472) and by the K427N mutation (Figure 3b). Interestingly, these features are conserved in human-infecting CoVs but not in sequences derived from animals. It is noteworthy that nearly all the inferred ancestral states had high support values (likelihood > 0.50; Supplementary Figure 4).

### Conserved mutations in SARS-CoV and SARS-CoV-2 RBD

Among the evolutionary changes identified in the RBD-specific SARS-CoV/SARS-CoV-2 clade (summarized in Figures 3b and 4a), residues 427N and 436Y are noteworthy for being conserved in SARS-CoV, SARS-CoV-2 and in other human-infecting strains but not in CoVs that only infect bats. Both mutations are in the short helix (residues 427-436) of the Spike protein, which lies at the interface between hACE2 and the Spike protein. The residue at position 436 forms hydrogen bonds with 38D and 42Q in the hACE2 structure (Figure 4d), likely contributing to the stability of the Spike-hACE2 complex; this interaction is known to be disrupted by the Y436F mutation^25^ found in RaTG13. A second mutation, K427N, is predicted to disrupt the RBD short helix, further reducing complex stability (Figure 4c). Both 436Y and 427N, are found in all SARS-CoV-2 isolates sequenced to date and also in viruses from other hosts, including civets (*Paguma larvata*) (Supplementary Table 1c) and the pangolin sequence collected in Guangdong. A mouse-adapted SARS virus also exhibits a mutation at position 436 (Y436H) that enhances the replication and pathogenesis in mice^26,27^, suggesting that this change may have affect host tropism. It is noteworthy that the 427N mutation is also found in strains not involved in the recombination event (KY770859 and KJ473816), suggesting that asparagine appeared in their most recent common ancestor through point mutation. Our ancestral state reconstructions (see below) also suggest that 427N has independently appeared two other times in bat SARS-like CoVs, but only at external branches (sequences JX993988 and JX993987; Figure 3a). Other bat isolates having the 427N allele, such as Rs7327, Rs4874 and Rs4231, are known to co-opt hACE2^28^, further reinforcing a role for 427N as an adaptive mutation involved in virus binding to hACE2.

**Fig. 4.**
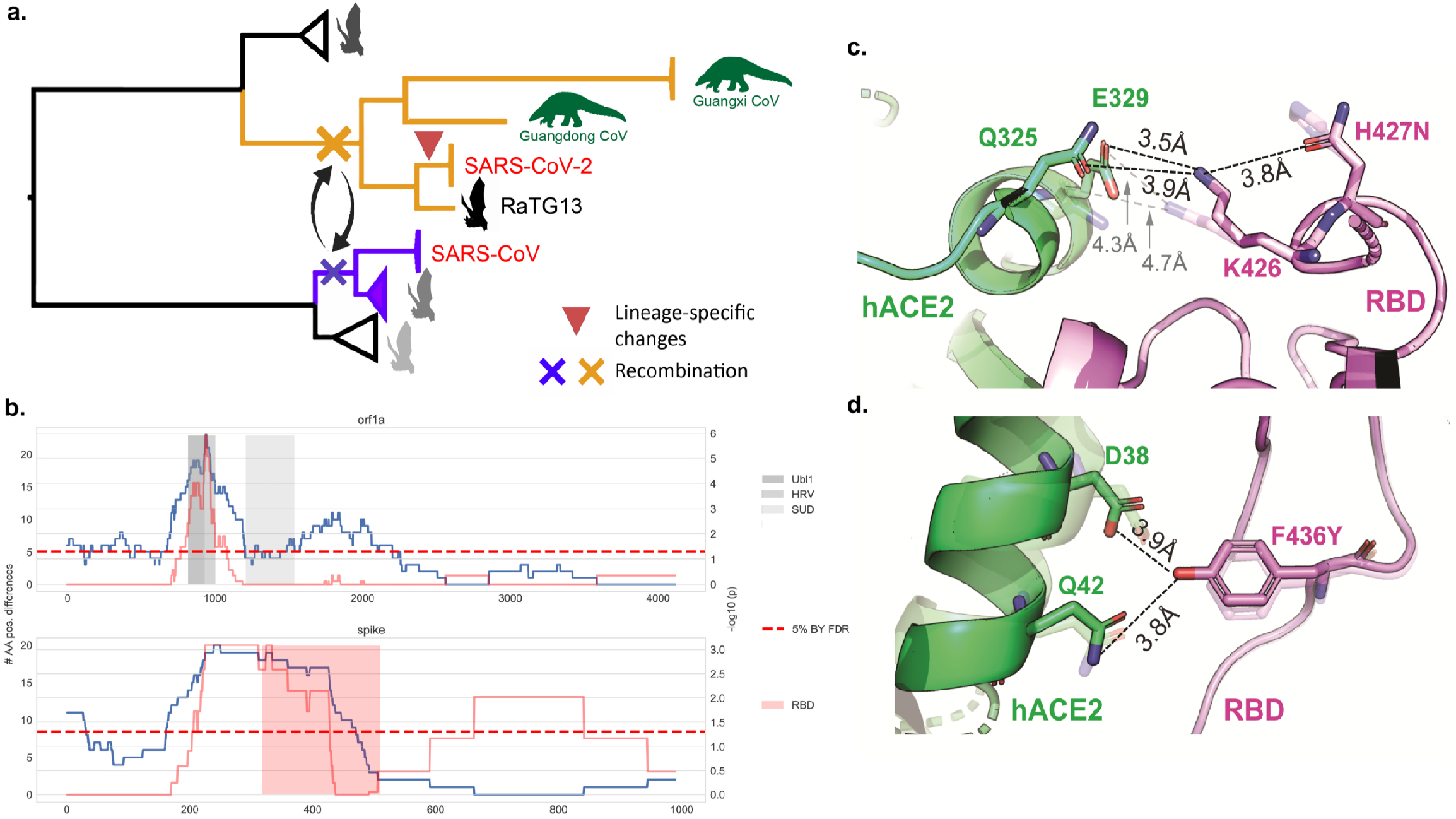
Evolutionary events preceding the emergence of SARS-CoV-2, and functional impact of amino acids 427N and 436Y. **a.** Phylogenetic representation summarizing the evolutionary events that likely led to the emergence of SARS-CoV-2: hit 1) Recombination of the RBD of the Spike protein involving lineages ancestral to SARS-CoV-2, RaTG13 and pangolin sequences (red cross) and SARS-CoV (blue cross); hit 2) SARS-CoV-2 accumulated four nonsynonymous mutations in RBD since its divergence from the MRCA that it shares with RaTG13 and the pangolin CoVs. **b.** Sliding window analysis (length 267 aa) identifies specific regions of SARS-CoV-2 with high divergence from the RaTG13 bat virus in the RBD of Spike (including 427N and 436Y), as well as in the Ubl1, HRV and SUD domains of nsp3 (non-structural protein 3) within the orf1a polyprotein. **c.** Functional impact of amino acid 427N in the SARS-CoV-2 Spike protein. Interaction between the human ACE2 receptor (green) and the spike protein (pink) based on SARS-CoV-2 (PDB accession code: 6LZG), highlighting the short helix 427-436 that lies at the interface of the Spike-ACE2 interaction. Dashed lines indicate hydrogen bonds between residues. The configuration shown with higher transparency is that of RaTG13 RBD interaction. **d.** Functional impact of amino acid 436Y.

### Recent substitutions in SARS-CoV-2 RBD

We compared the SARS-CoV-2 sequence to the bat CoV nearest in sequence across the entire genome, namely RaTG13. The distributions of nonsynonymous and 4-fold degenerate site changes between SARS-CoV-2 and RaTG13 across the viral genome revealed two regions with significant enrichment of nonsynonymous changes (adjusted *p*-val. < 10^-5^ and *p*-val. < 10^-3^ for the first and second regions respectively, binomial test on sliding windows of 267 amino acids) (Figure 4b). The first region, analyzed using sliding windows that covered nucleotide positions 801 to 1067 in the orf1a gene, spans the non-structural proteins (nsp) 2 and 3 that were previously reported to accumulate a high number of mutations between bat and SARS CoVs.^29^ These region include the ubiquitin-like domain 1, a glutamic acid-rich hypervariable region, and the SARS-unique domain of nsp3^30^ that is critical to replication and transcription^31,32^. The second region with high divergence from RaTG13 contained 27 substitutions in the Spike protein, of which 20 were located in the RBD (Supplementary Table 2). No significant enrichment was observed for mutations at 4-fold degenerate sites (Supplementary Figure 5).

Through ancestral state reconstruction, we also identified four nonsynonymous substitutions in the RBD that were specific to the lineage leading to SARS-CoV-2 (Supplementary Table 3). The four amino acid changes in RBD, with respect to their respective ancestral states, are T333R, T359A, H484Q and N505H (where the reference allele is indicated on the left-hand side). Interestingly, residue 484Q has already been reported to interact directly with hACE2^33^.

### Increased binding affinity of RBD to hACE2 is associated with the ancestral recombination

We performed structural modeling of the RBD and hACE2 binding (using trRosetta^34^ and the Rosetta energy minimization scripts^35^) to infer how the binding affinity of Sarbecoviruses to the human receptor has changed with evolution, focusing on the clade implicated in the ancestral RBD recombination at positions 22614-23032 (see above; this clade includes SARS-CoV, SARS-CoV-2, pangolin CoVs from Guangdong and RaTG13). Using the reconstructed ancestral states, we estimated changes in binding affinity for hACE2 along one evolutionary trajectory leading to SARS-CoV and a second trajectory leading to SARS-CoV-2. We observed that both the common ancestor for the whole RBD recombination clade (−36.91 kcal/mol), and that of SARS-CoV and SARS-CoV2 (−34.58 kcal/mol) had higher binding affinity than the ancestor preceding this recombination clade, or than an external bat-SL-CoV (−16.32 kcal/mol) (Figure 5). Subsequent vertical evolution resulted in more modest increases in binding affinity of MRCAs leading to SARS-CoV: −40.89 kcal/mol for the MRCA of SARS-CoV and its closest bat CoVs (step ‘4A’, Figure 5); and −39.95 kcal/mol for SARS-CoV (step ‘4B’, Figure 5). A similar trend (−47.38 kcal/mol overall) was observed along the evolutionary trajectory leading to SARS-CoV-2 (steps ‘3A’ and ‘3B-S’, Figure 5). Indeed, the most recent common ancestor of SARS-CoV-2, pangolin CoVs from Guangdong and RaTG13 (‘3A’, Figure 5) had an increased binding affinity by more than 20 kcal/mol as compared to the ancestor prior to the RBD recombination event. Both the pangolin CoV and RaTG13 had a lower affinity than their own common ancestor with SARS-CoV-2. The predicted affinity of SARS-CoV-2 (‘3B-S’) is the highest, recapitulating observations that SARS-CoV-2 has a higher binding affinity to hACE2 than SARS-CoV (Figure 5). We conclude that recombination in the RBD substantially increased the hACE2 affinity of an ancestral lineage to SARS-CoV-2.

**Fig. 5.**
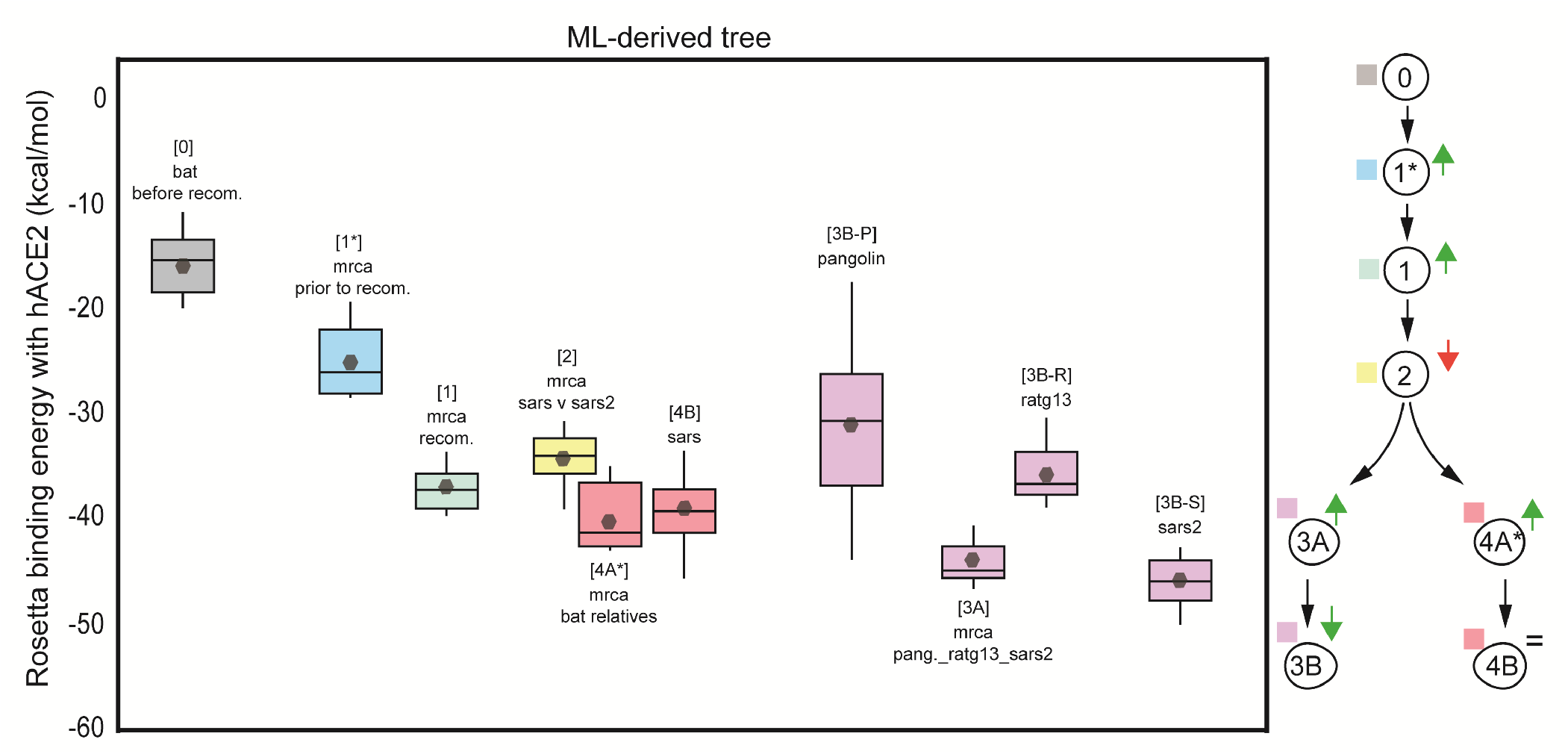
The ancestral recombination event at RBD involving SARS-CoV/SARS-CoV-2 is associated with increased binding affinity to hACE2. Boxplots represent the distribution of the binding energies of the RBD of each viral strain to hACE2, as inferred by Rosetta. Viral strains (and the analyzed MRCAs) have been labelled with numbers in a hierarchical order, as follows. Outgroup sequence: 0, MRCA from which the MRCA of the recombination cluster derives: 1, MRCA of the recombination event: 2, MRCA of SARS-CoV and SARS-CoV-2 lineages: 1, MRCA of SARS and its bat-SL-CoV relatives: 4A, SARS-CoV: 4B, MRCA of RaTG13, Pangolin-CoV and SARS-CoV-2: 3A, Pangolin-CoV: 3B-P, RaTG13: 3B-R, SARS-CoV-2: 3B-S. The diagram on the right summarizes the progressive increase in binding affinity along the evolutionary trajectories leading to SARS-CoV and SARS-CoV-2. All strains involved in the SARS/SARS-CoV-2 recombination event, including their MRCA, exhibit higher binding affinity (lower binding energy) than the bat SARS-like CoV used as outgroup (MG772933, ‘0’). Binding affinity increased further along the evolution of human-infecting Sarbecoviruses (SARS-CoV, SARS-CoV-2). The highest binding affinity among all strains analyzed is found in SARS-CoV-2 (‘3B-S’) and its MRCA shared with Pangolin-CoV and RaTG13 (‘3A’).

We also assessed the effects of the SARS-CoV-2 lineage-specific point mutations on binding affinity by repeating the Rosetta analyses of SARS-CoV-2 RBD, but reverting alleles at positions 333, 359, 484 and 505 to their ancestral states individually and also in all possible combinations. None of them had an effect on binding affinity to hACE2 with the exception of Q484H, which had a significant effect on increasing hACE2 binding affinity (Figure 6A). We performed ten independent Rosetta trajectories per mutant to determine the median scores for binding to hACE2 when all nineteen amino acid point mutations are made at residues 333, 359, 484 and 505. Our results are in line with those reported through independent experimental assays^36^ (Supplementary Figure 6).

**Fig. 6.**
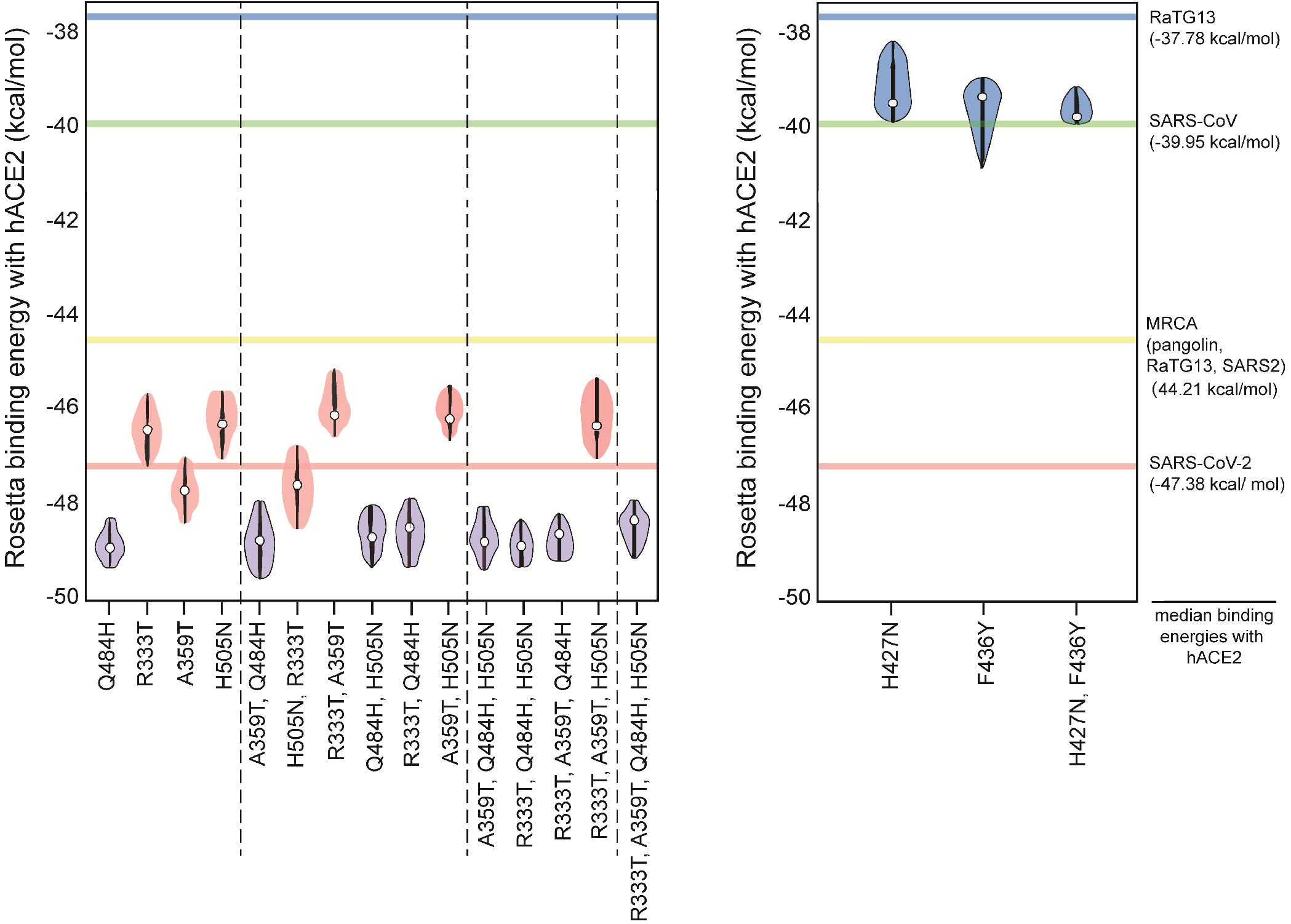
The effects of specific alleles on RBD binding affinity to hACE2. **a.** Change in the binding energy of SARS-CoV-2 RBD to hACE2 caused by the reverse mutation of each SARS-CoV-2 lineagespecific allele to its ancestral state. Binding energy was assessed by considering each mutation individually as well as all possible combinations among the four different SARS-CoV-2 lineage-specific amino acids. **b.** Binding energy of RaTG13 RBD to hACE2 after mutating positions 427 and 436 (either individually or both together) to the SARS-CoV/SARS-CoV-2 alleles.

Finally, we assessed the relevance of 427N and 336Y on binding to hACE2 (Figure 4c, d). Given that RaTG13 appears to have lost these two alleles (having instead 427H and 336F), we modelled binding after incorporating the H427N and F426Y mutations individually and together. Individually, these alleles had a significant effect on improving binding affinity of RaTG13 to hACE2. However, the biggest effect was found when they were present in the same RBD sequence, raising the affinity of RaTG13 and giving it an affinity to bind hACE2 similar to that of SARS-CoV (Figure 6B).

## Discussion

In this work, we analyze the evolution of SARS-CoV-2 and its closest relatives, with a focus on the RBD region of the Spike protein, as a means to better understand viral tropism. It has been hypothesized previously that recombination^15,37^ and rapid evolution has occurred in bat, civet and human SARS-CoVs^29^. However, previous descriptions of recombination in the Spike protein were purely observational^15^. In contrast, we use statistical methods to show that recombination events preferentially affect the RBD region in the Spike gene, both at the level of the genus (betacoronavirus) and within individual species (such as MERS-CoV). This enrichment for recombination found at Spike is in agreement with recently published work^38^. Our analyses suggest that the evolutionary history of the RBD from human-infecting CoVs is characterized by an ancestral recombination event that would involve the ancestors of SARS-CoV and SARS-CoV-2. Unlike in the rest of the genome, in the RBD SARS-CoV and its closest related strains belong to a well-supported cluster together with SARS-CoV-2, RaTG13 and the pangolin CoVs. Interestingly, RmYN02 lies far away from the SARS/SARS-CoV-2 cluster in the RBD, suggesting that this bat strain has undergone some further recombination event at Spike. This further recombination involving RmYN02 is in agreement with recent publications^39^.

Importantly, the ancestral recombination event reported here, involving ancestors of SARS and SARS-CoV-2, had a significant impact on the hACE2-binding affinity of the RBD, as did the occurrence of subsequent key amino acid changes in RBD. A main caveat of our analysis is that we cannot unveil which CoV strains are the recombinant or the parental ones, likely because of the ancestry of the event and the potential effect of undersampling. However, our randomization tests comparing the RBD to the whole-genome tree revealed that SARS-CoV together with SARS-CoV-2 and its closest strains contributed the most to the dissimilarity between the two phylogenetic trees. This supports a recombination event involving ancestors of these two human-infecting CoV strains.

Recent work has suggested that other recombinations may have occurred involving SARS-CoV-2, pangolin CoV and RaTG13 in the Spike gene. Such an event was proposed to explain the drop in nucleotide similarity between SARS-CoV-2 and RaTG13^18^. However, other papers have argued against such a recombination event and suggested that if any recombination event could explain such a drop in similarity, then it would only involve RaTG13 and an unsampled (unknown) parent^19^.

Modeling Spike-hACE2 binding using trRosetta, the current state-of-the-art in rapid and reliable *de novo* prediction of protein structure, shows that the protein generated by the proposed recombination event involving ancestral strains of SARS-CoV/SARS-CoV-2 increased affinity to hACE2. This is exemplified by the significantly higher binding affinity displayed by the most recent common ancestor (MRCA) of the clade implicated in the ancestral RBD recombination, as compared to sequences external to this clade. Structural modeling of different RBD sequences in the reconstructed evolution of SARS-CoV and SARS-CoV-2 shows that each lineage enhanced affinity for hACE2 binding. Strikingly, the predicted affinity of the inferred common ancestor for RaTG13, pangolin CoV and SARS-CoV-2 is predicted to have a high affinity to hACE2, (5.66 kcal/mol) higher than that of SARS-CoV but 1.76 kcal/mol lower than that of SARS-CoV-2, which has the highest affinity of all the sequences that we studied. These findings suggest that the postulated common ancestor between RaTG13, SARS-CoV-2 and pangolin CoV had the ability to infect humans. These ancestors must have circulated in animal populations prior to 2013, when RaTG13 was collected, suggesting that viruses infectious to humans may have been present in the wild for decades before a zoonotic jump and human to human transmission. In line with our results, recent experimental analyses have revealed that bat-SL-CoVs closely related to SARS-CoV (strains WIV1 and SHC014) efficiently replicate in human cells without undergoing any adaptive process^40^.

The presence of such ancestral strains prior to SARS-CoV-2 emergence is supported by the tMRCA inferences made by other methods, which have suggested that the split between RaTG13 and SARS-CoV-2 could indeed have happened decades ago^19,41^.

Our results suggest that recombination was a key factor in the emergence of Sarbecoviruses into humans, resolving the debate about whether recombination or convergent evolution were involved in the emergence of SARS-CoV-2^7^. We also found that Spike protein positions 427N and 436Y are retained in human Sarbecoviruses but are not conserved among non-human strains. Residue 436Y in the Spike is predicted to be part of the RBD-ACE2 interface, while 427N is contiguous to 426K, which is a key residue for establishing strong electrostatic stabilization of hACE2 residues 325Q and 329E^33,9^ (Figure 4c).

Previous reports focused on the differences between SARS-CoV-2 and alphacoronaviruses or on sites that differ between SARS-CoV-2 and SARS-CoV^42^. Others, have suggested through dN/dS analyses that the lineage leading to SARS-CoV-2 and its closest strains (RaTG13, RmYN02, pangolin CoVs) would have undergone in mammals (e.g. bats) an adaptive process facilitating infection in humans^39^. However, no analyses were performed to trace and compare the evolution of the receptor binding affinity in ancestors that led to SARS-CoV-2. In contrast, in this work we identify potentially important loci from evolutionary analysis and inference of ancestral alleles. Modeling suggests that SARS-CoV-2 binds significantly more tightly to the human receptor than SARS-CoV. On the other hand, pangolin CoV from Guangdong and RaTG13 differentiated outside human hosts, decreasing their binding affinity towards hACE2. In this way, we have quantitatively assessed the binding affinity of SARS-CoV, SARS-CoV-2 and its closest strains and ancestors and validated *in silico* the role of positions 427 and 426.

Previous analyses have also reported other unique features of SARS-CoV-2 at the Spike protein. For instance, the insertion of a four-residue polybasic cleavage site that is only present in SARS-CoV-2 and RmYN02^6^ among the Sarbecoviruses; this site lies between Spike positions 666-667 (in SARS-CoV NC_004718 coordinates). The presence of polybasic cleavage sites has been associated in viruses such as Influenza with high pathogenicity^9,43^, and experiments that introduce such sites into the S1-S2 junction, between Spike codon positions 667 and 668 (SARS-CoV-1 coordinates) of human SARS-CoV result in an increase in cell-cell fusion without affecting viral entry^42,44^.

In conclusion, evolutionary analyses and structural modeling show that the evolutionary processes giving rise to SARS-CoV-2 included a recombination involving ancestors of SARS-CoV and SARS-CoV-2, followed by the accumulation of point mutations in the Spike protein. Both the ancestral recombination event and the point mutations, which differ between SARS-CoV and SARS-CoV-2, resulted in progressively tighter binding to hACE2. It appears that ancestors to SARS-CoV-2, with the ability to bind tightly to hACE2 and thus potentially infect humans, may have been circulating in the wild for decades prior to making the jump to humans and causing pandemic disease. These results exemplify the importance of combining evolutionary analyses with protein structure and binding affinity predictions in order to assess the host-switching potential of animal-infecting viruses based on the genetic changes that have accumulated along their evolution.

## Author Contributions

J.P., I.F. and R.R. designed the study and prepared the manuscript. J.P. and I.F. performed computational analyses. R.C., C.M., M.A., and P.K.S. designed and performed protein structure analyses. All authors contributed to writing the manuscript.

## Acknowledgements

We thank GISAID and all the laboratories where the data used in this study was collected and processed (Supplementary Table 4). We would also like to thank Karen Gomez and Andrew Chen for their help editing the manuscript, and Zixuan Wang for her help on the figures. The icons for the host species in our figures were created with BioRender. Finally, this work has been funded by NIH grants R01 GM117591 and U54-CA225088 and by DARPA/DOD grant W911NF-14-1-0397.

## Online Methods

### Sample collection: SARS-CoV-2 and SARS/SARS-like-CoVs

A set of 71 genome sequences derived from SARS-CoV-2 (which represent all genome availability at GISAID on February 7, 2020; gisaid.org) was analyzed together with its closest animal-infecting relative, RaTG13 (accession number MN996532), sequences sampled from pangolins (n=5) and other genome sequences from human SARS-CoV (n=72) and bat SARS-like CoV (n=39), publicly available in Genbank (ncbi.nlm.nih.gov/genbank/) (Supplementary Table 1d). Alignment was performed either at genome wide nucleotide level or at the Spike CDS (at amino acid level)independently with MAFFTv7 (“auto” strategy)^45^. Seven recombination detection methods implemented in the RDP4 software package (RDP, Geneconv, Bootscan, Maxchi, Chimaera, SiScan, 3seq)^22^ were used to detect evidence of recombination with default parameters (*p*-value = 0.05, Bonferroni corrected), and depict the distribution of recombination events, in different CoV alignments:

1. The following selection of viral strains was used in order to find breakpoints involving SARS-CoV-2: KF636752 (bat), NC_004718 (human SARS), DQ071615 (bat SARS-like CoV), DQ412043 (bat SARS-like CoV), MG772933 (bat SARS-like CoV, MN996532 (RaTG13), NC_045512 (SARS-CoV-2).
2. A MERS-CoV genome alignment (n= 381; n= 170 human, n= 209 camel, and 2 bat sequences).
3. A betacoronavirus alignment (n=45 sequences, covering the genus diversity as in *Lu, R. et al., 2020*^21^).

Any recombination event detected was further considered only if it met two additional criteria:

i. The potential recombination event should be phylogenetically informative. This was evaluated by means of likelihood-mapping analyses of 1000 random tree quartets in Tree-Puzzle^46^: those recombinant segments leading to >30% of unresolved quartets (those with star-like signal) were no further considered.
ii. The tree topology derived from such recombinant segment should be significantly different from that obtained from the rest of the genome. For each event, we compared both ML trees (obtained with PhyML^47^ with a GTR + GAMMA 4 CAT substitution model, thus accounting for among-site rate variation) by means of Expected Likelihod Weight (ELW) and Shimodaira-Hasegawa (SH) tree topology comparison tests implemented in Tree-Puzzle, and we only considered those events that passed both tests.

### Phylogenetic analysis

The evolutionary relationships between SARS-CoV-2 and other SARS/SARS-like CoVs was inferred from genome alignment using PhyML (GTR + GAMMA 4CAT)^47^. The same program and model were used to reconstruct the phylogenetic tree of the (potentially recombinant) RBD.

In order to get further support that the reported recombination event at RBD involved ancestors of SARS-CoV and SARS-CoV-2, we assessed the impact of the SARS-CoV/SARS-CoV-2 clade (see Figure 3a) on the divergence between the maximum likelihood phylogeny of the whole viral genome and that of the recombining RBD segment (denoted below by T1 and T2 respectively). We performed this assessment by means of a randomization test. Specifically, we first measured the distance between T1 and T2 in the Villera–Holmes–Vogtmann (BHV) metric^24^, which compares the tree topologies and internal branch lengths of phylogenetic trees, using the GTP algorithm^48^. We note that the BHV metric provides a natural method to assess the dissimilarly (or “divergence”) between two phylogenetic trees because it incorporates both topological differences and the edge lengths, unlike the classical Robinson–Foulds metric which only accounts for splits present in exactly one of the two input trees^49^. Second of all, we compared the BHV distance between T1 and T2 with the distance between the corresponding truncated trees (“truncated T1” and “truncated T2”), obtained by pruning all the sequences (or nodes) that comprised the SARS-CoV/SARS-CoV-2 clade (20 sequences in all). Finally, the difference between these two BHV distances quantifies the impact of the recombining SARS-CoV/SARS-CoV-2 clade to the overall divergence between the whole-genome phylogeny and the RBD-specific phylogeny.

Differences of BHV distances (i.e. full-tree BHV distance minus truncated BHV distance) was also calculated for random node samples of varying sizes, sampling from 1 to 25 nodes at a time (in total, 23780 random sets of sequences were analyzed, with 1000 samples per size of the sampled subset). A *p*-value for the significance of the SARS-CoV/SARS-CoV-2 clade (composed of 20 SARS-CoV-2 related sequences, see Figure 3a) was derived as the proportion of node samples for which the difference

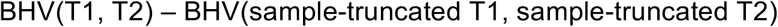

was strictly greater than the corresponding difference calculated for the SARS-CoV/SARS-CoV-2 clade. A low *p*-value for the SARS-CoV/SARS-CoV-2 clade (*p*-val. < 0.0001) thus indicates that the 20 sequences which compose this clade are contributing significantly to the divergence between the wholegenome and the RBD phylogenies of Sarbecoviruses. Without identifying specifically the recombinant sequence, this analysis nevertheless supports our hypothesis that ancestors of SARS-CoV and SARS-CoV-2 were directly implicated in the ancestral RBD recombination reported in this work at coordinates 22614-23032. We obtained similar results when we computed differences between BHV distances normalized alternatively: either by the total number of sequences remaining in the pruned trees; or by re-weighing tree edge-lengths so that the squared sum of internal branch lengths always equals 1.

### Ancestral state reconstruction and RBD-hACE2 binding affinity analyses

The amino acid changes that occurred in the Spike gene along the evolution Sarbecoviruses were traced using a Maximum Likelihood state reconstruction approach implemented in the Ape R package^50^. As input, the ML tree derived from the RBD recombinant region was used. ML is used to fit a markov model of discrete character evolution where all transitions between characters had the same probability (equal rates model, ER). It asks what is the most likely state for each node in the tree, integrating over all the possible states, over all the other nodes in the tree.

The binding affinity towards hACE2 of the reconstructed RBD from different ancestors (MRCA of the event, MRCA between SARS-CoV and SARS-CoV-2, MRCA of SARS-CoV and its closest bat SARS-like-CoV and MRCA of Pangolin-Guangdong, RaTG13 and SARS-CoV-2) was then evaluated, and compared with the affinity of MG772933 (sequence external to the reported ancestral recombination event), SARS-CoV, pangolin-Guagdong, RaTG13 and SARS-CoV-2. To measure RBD binding affinity towards hACE2, the nine RBD-hACE2 complexes were built using the experimentally determined atomic coordinates of the SARS-CoV-2 RBD-hACE2 complex (PDB accession 6lzg^51^) as a reference. To this end, we first used the trRosetta^34^ structure prediction algorithm to generate seven of the nine RBD structures (except SARS-CoV and SARS-CoV-2, which had experimentally reported crystal structures in complex with hACE2). These complexes were subsequently relaxed along ten independent trajectories each, using Rosetta quasi-Newton FastRelax energy-minimization scripts (with inexact linear search Broyden-Fletcher-Goldfarb-Shanno -BFGS- update method^52^). The average and median binding energies (in kcal/mol), and the number of interface hydrogen bonds were then computed based on these trajectories for each of the nine RBD-hACE2 complexes.

We also assessed the effect of SARS-CoV-2 lineage specific RBD mutations H484Q, T333R, T359A and N505H (those detected in the ancestral state analyses) on the binding affinity of SARS-CoV-2. We used the RBD sequence of SARS-CoV-2 and reverted those positions to their ancestral state, either individually or in combination, using the RosettaMP application. Next, we performed a rotamer repacking protocol to relax the side chain conformations of all RBD residues within 10Å of the mutated residue to alleviate steric clashes and establish stabilizing contacts. After docking the mutated RBDs with hACE2, the resultant complexes were fed through the aforementioned Rosetta energy-minimization pipeline. Ultimately, the binding energy calculations were used to infer the effect of all possible combinations of point mutations of SARS-CoV-2 on hACE2 binding. We also assessed the effect of the human-associated alleles 427N and 436Y on binding affinity to hACE2. These two alleles are fixed in human-infecting Sarbecoviruses, but not in non-human isolates (and are not present in RaTG13). For this reason, we mutated RaTG13 RBD into H427N and F436Y and assessed whether introducing these human alleles leads to an increase in binding affinity of RaTG13 RBD. For this, we followed the same procedure as in the aforementioned SARS-CoV-2 specific mutations.

### Statistical analysis

Sliding window analysis was performed in order to test for enrichment of recombination breakpoints (including both start and end breakpoints) along the viral genome in the following settings: 1) all beta-CoV recombinations; 2) recombinations within non-human lineages for beta-CoV; 3) all MERS-CoV recombinations; and 4) both human-specific and non-human MERS-CoV lineage recombinations separately. There were too few human-specific recombinations in beta-CoV for in-depth analysis. For beta-CoV analyses, the SARS-CoV genomic coordinates were used as reference (accession NC_004718), whereas for MERS CoVs, we used a MERS-CoV sequence (accession NC_019843) as reference. Windows of 800 nucleotides were selected and binomial tests for the number of breakpoints in each window were performed under the null hypothesis that recombination breakpoints are distributed uniformly along the genome. Given the co-dependence structure of our statistical tests, adjustments were performed using the Benjamini-Yekutieli (BY) procedure^53^ which provides a conservative multiple hypothesis correction that applies in arbitrary dependence conditions. For statistical significance, we chose 5% BY false discovery rate (FDR). Our discoveries are valid with different choices of window length, provided the window length is sensitive to the scale CoV proteins and the length of specific domains such as the RBD in the Spike gene.

We used the same sliding window approach to test for enrichment of gene-specific nonsynonymous as well as synonymous differences between SARS-CoV-2 and the bat virus RaTG13. For consistency, we selected 267 length windows of amino acids (corresponding to approximately 800 nucleotides) and performed *p*-value correction using the same procedure.

