## Supplementary Figures for "Recombination and lineage-specific mutations linked to the emergence of SARS-CoV-2"

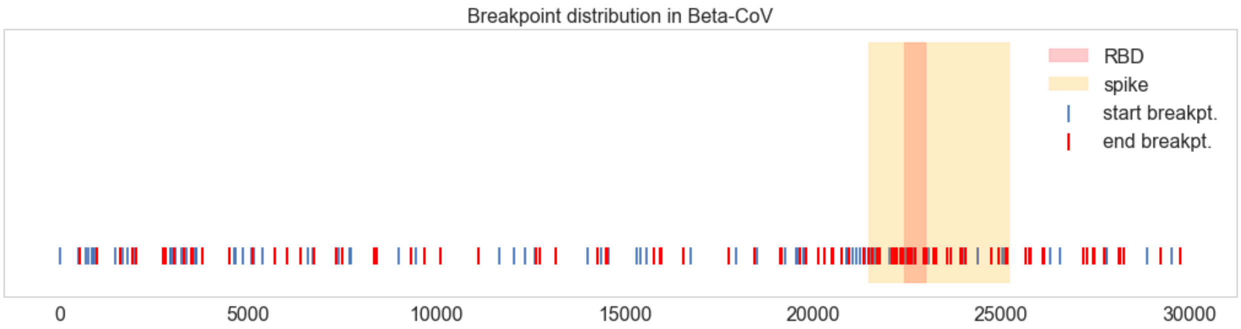


**Supplementary Figure 1a | Recombination breakpoints in betacoronaviruses.** Start (blue) and end (red) breakpoints from all of the 103 detected recombination events in betacoronaviruses are shown, highlighting the Spike protein and the RBD.

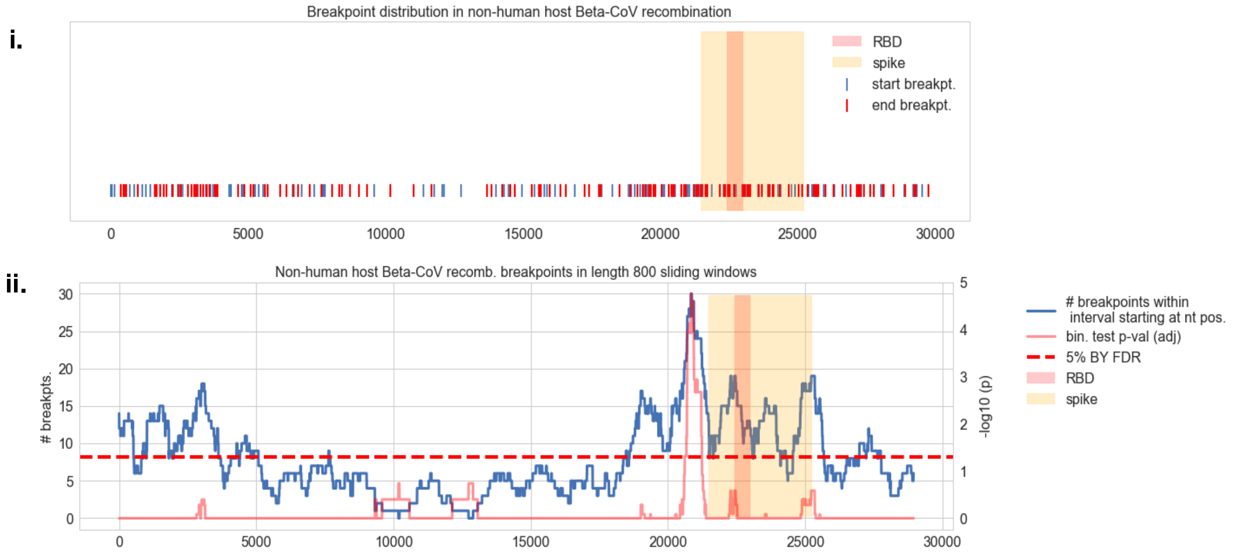


**Supplementary Figure 1b | Recombination breakpoints in betacoronaviruses (non-human host only). i.** Distribution of start and end breakpoints recombination events detected in betacoronaviruses infecting non-human hosts. The Spike protein and the RBD are highlighted. **ii.** Sliding window analysis shows the distribution of recombination breakpoints (either start or end) in 800 nucleotide (nt) length windows upstream (namely, in the 5' to 3' direction) of every nt position along the viral genome (blue curve). BY-adjusted *p*-values are shown in red.

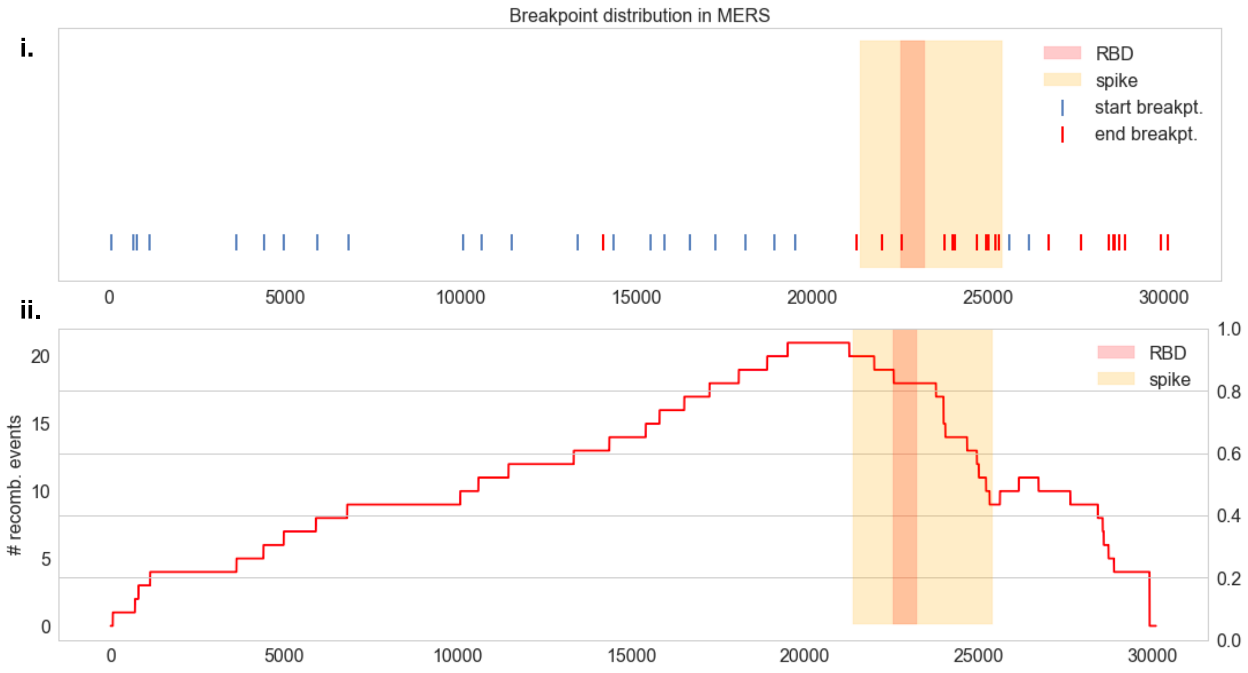


**Supplementary Figure 1c | Recombination breakpoints in MERS-CoV. i.** Distribution of start and end breakpoints from 24 recombination events detected from MERS-CoV isolates. **ii.** Number of recombination events covering each nt position of the MERS-CoV genome. The Spike gene, and the n-terminus in particular, overlaps with the majority of the detected recombination segments.

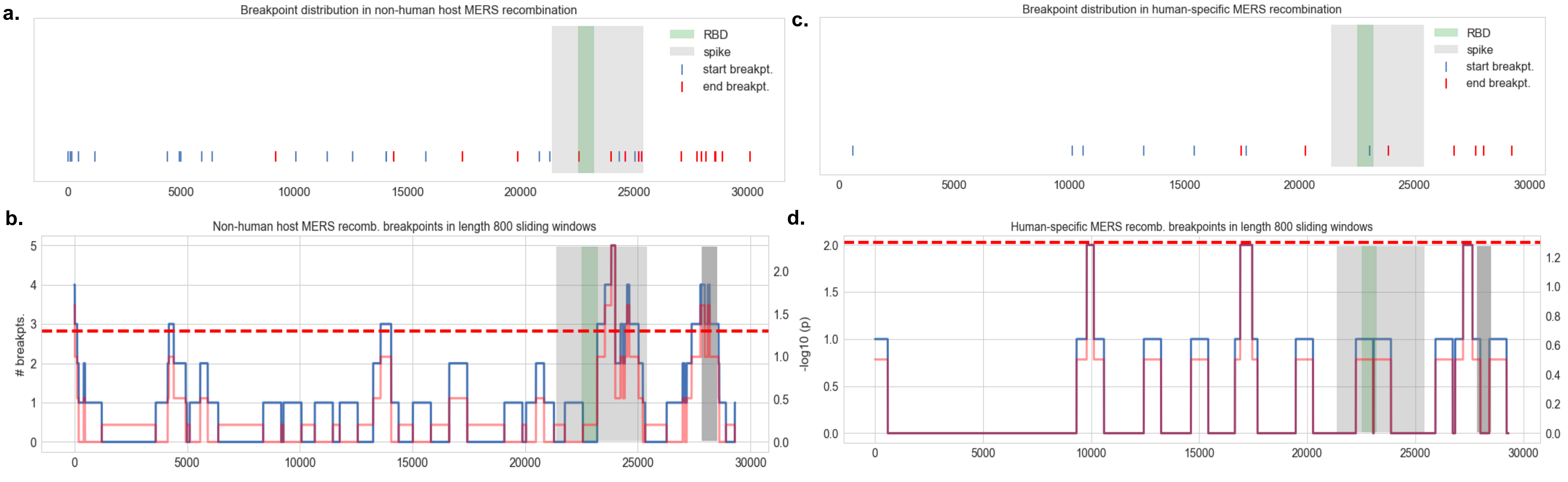


**Supplementary Figure 2 | Host-specific recombination breakpoints in MERS-CoV. a, c.** Distribution of start and end recombination breakpoints in MERS-CoV isolates from non-human hosts (a) and human hosts (c). **b, d.** Sliding window analysis: 800 nt length windows for non-human (b) and human-specific recombination events (d); recombination breakpoint count in each interval shown in blue; binomial test nominal p-values in red. Spike protein (light gray), RBD of Spike (green) and membrane protein (dark gray) are highlighted.

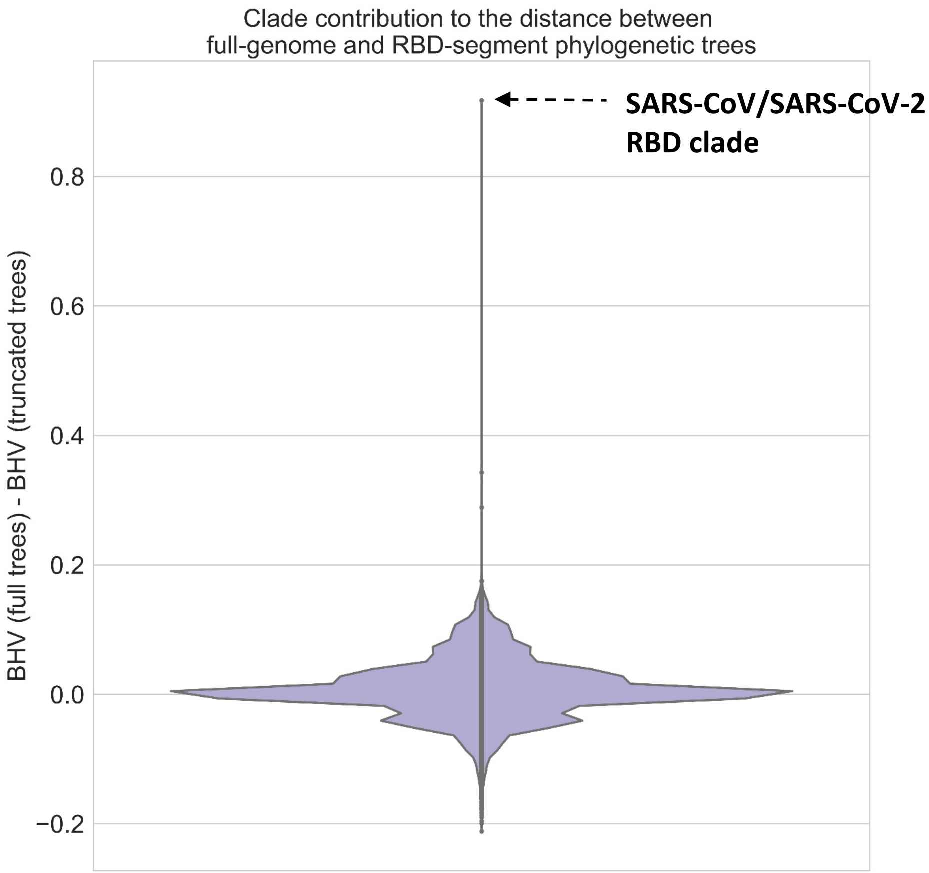


**Supplementary Figure 3 | SARS-CoV and SARS-CoV-2 lineages contribute significantly to genome vs RBD tree differences.** Violin plot showing the difference of BHV distances between the full-genome tree and the RBD-specific tree on the one hand, and the corresponding truncated trees on the other hand. Differences for 23780 random sets of sequences and clades are shown. The difference of BHV distances quantifies the contribution of the SARS-CoV/SARS-CoV-2 clade towards the overall divergence between the whole-genome and the RBD evolutionary histories, and this randomization test shows the probability that a randomly selected set of taxa contributes more towards the phylogenetic divergence of the full-genome and the RBD-specific trees than the SARS-CoV/SARS-CoV-2 clade. The SARS-CoV/SARS-CoV-2 clade ranks the highest among all the clades and random sets of taxa analyzed.

**Supplementary Figure 4 | Distribution of Likelihood values of the RBD amino acids reconstructed for the differences ancestors of interest.**

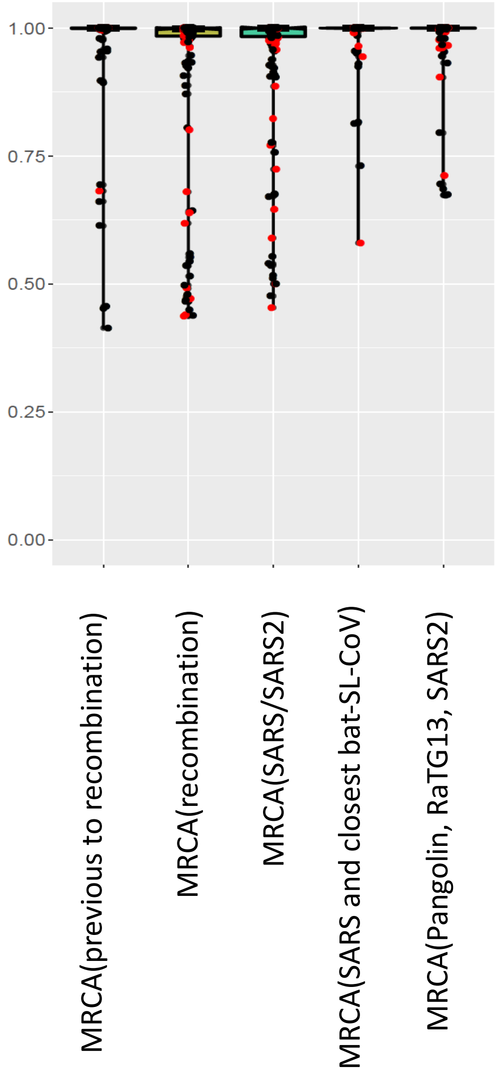

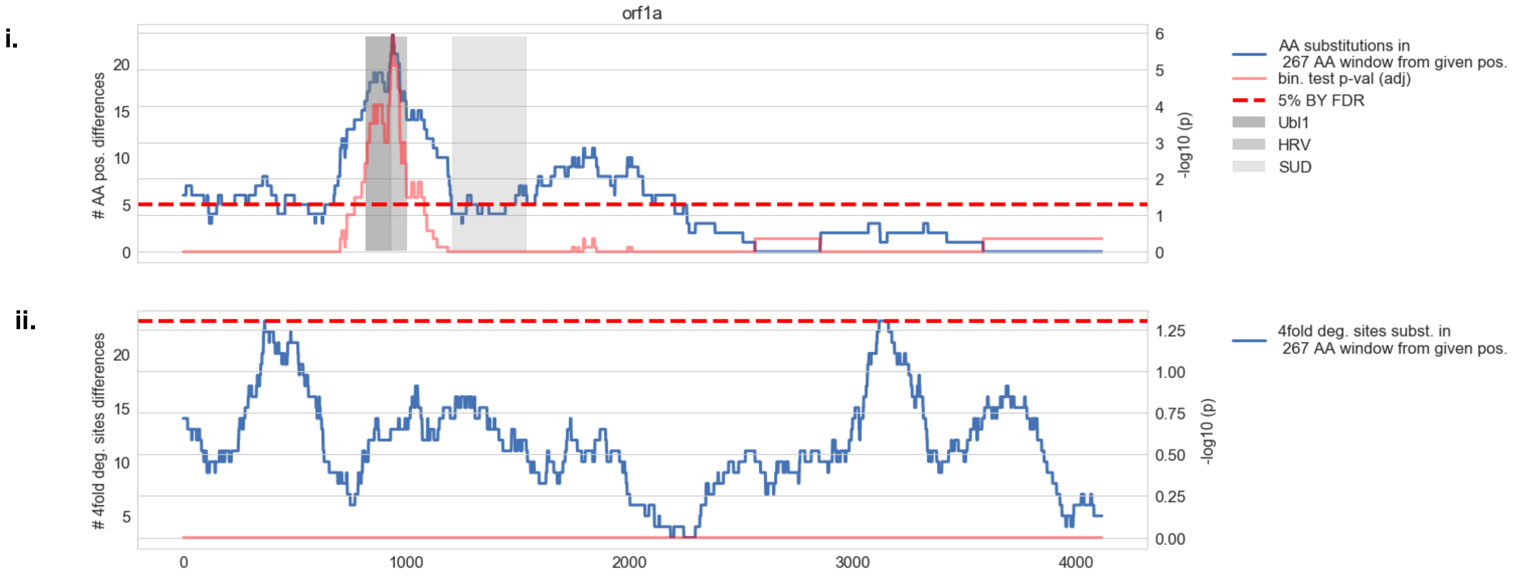


**Supplementary Figure 5a | Divergence between SARS-CoV-2 and the RaTG13 bat virus in orf1a. i.** Sliding window analysis of 267 aa length windows showing non-synonymous substitutions where the two strains differ (blue). BY-adjusted *p*-values for each window shown in red. Several domains of non-structural protein three are highlighted in gray: ubiquitin-like domain 1 (Ubl1), hypervariable region (HRV) and SARS-unique domain (SUD). **ii.** Sliding window analysis (267 aa length windows) containing 4-fold degenerate sites that differ between SARS-CoV-2 and RaTG13. No windows are statistically significant after BY *p*-value adjustment (red line).

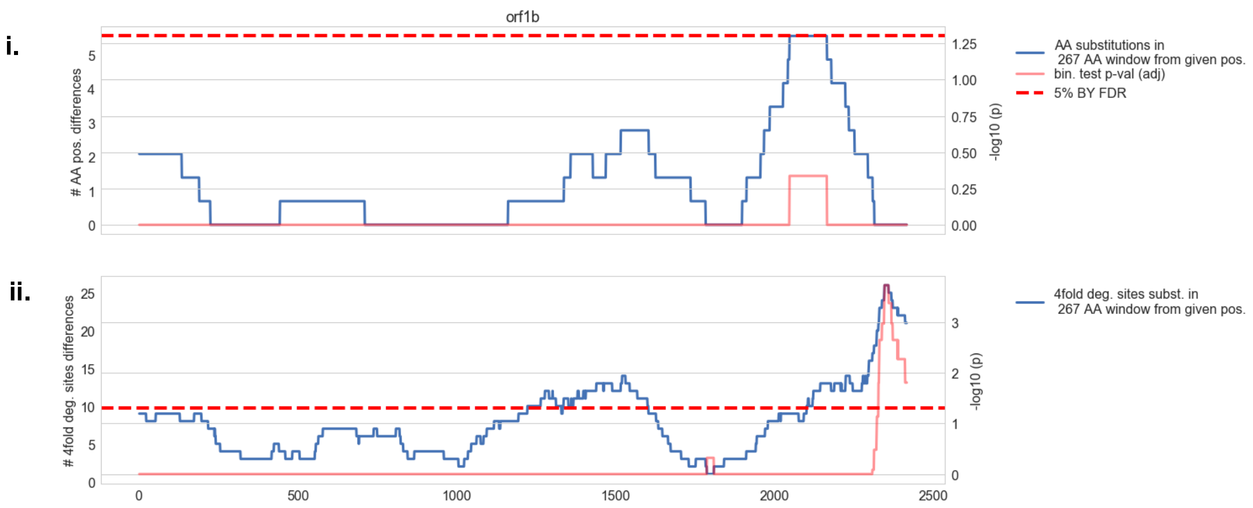


**Supplementary Figure 5b | Divergence between SARS-CoV-2 and the RaTG13 bat virus in orf1b. i.** Sliding window analysis of 267 aa length windows showing non-synonymous substitutions where the two strains differ (blue). BY-adjusted *p*-values for each window shown in red. **ii.** Sliding window analysis (267 aa length windows) containing 4-fold degenerate sites that differ between SARS-CoV-2 and RaTG13.

**i.**

**ii.**

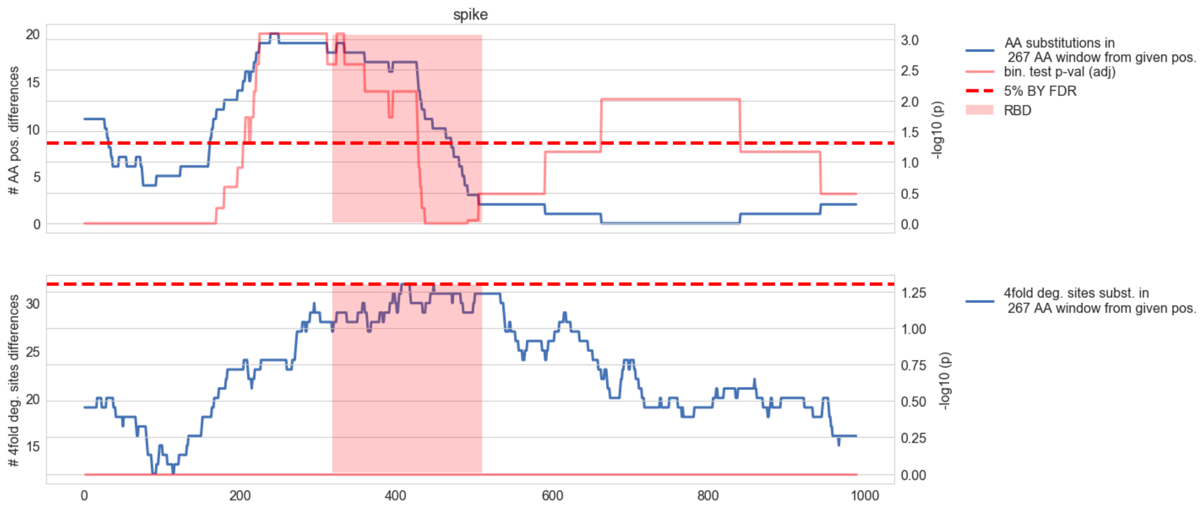


**Supplementary Figure 5c | Divergence between SARS-CoV-2 and the RaTG13 bat virus in Spike. i.** Sliding window analysis of 267 aa length windows showing non-synonymous substitutions where the two strains differ (blue). BY-adjusted *p*-values for each window shown in red. RBD is highlighted. **ii.** Sliding window analysis (267 aa length windows) containing 4-fold degenerate sites that differ between SARS-CoV-2 and RaTG13.

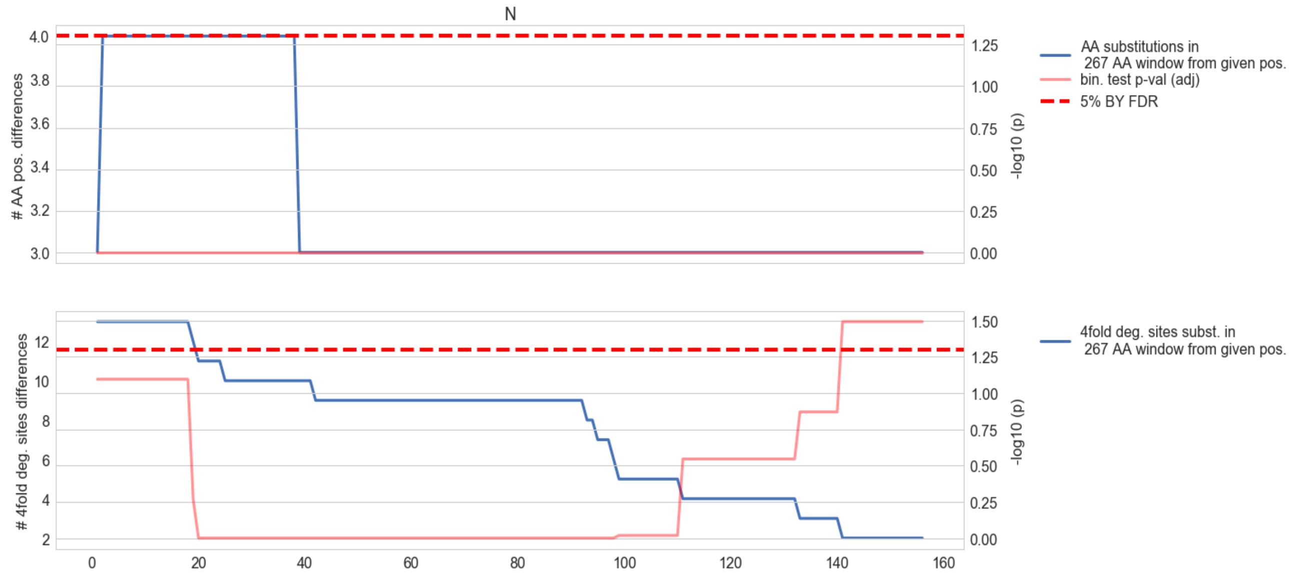


**Supplementary Figure 5d | Divergence between SARS-CoV-2 and the RaTG13 bat virus in the Nucleocapsid. i.** Sliding window analysis of 267 aa length windows showing non-synonymous substitutions where the two strains differ (blue). BY-adjusted *p*-values for each window shown in red. **ii.** Sliding window analysis (267 aa length windows) containing 4-fold degenerate sites that differ between SARS-CoV-2 and RaTG13.

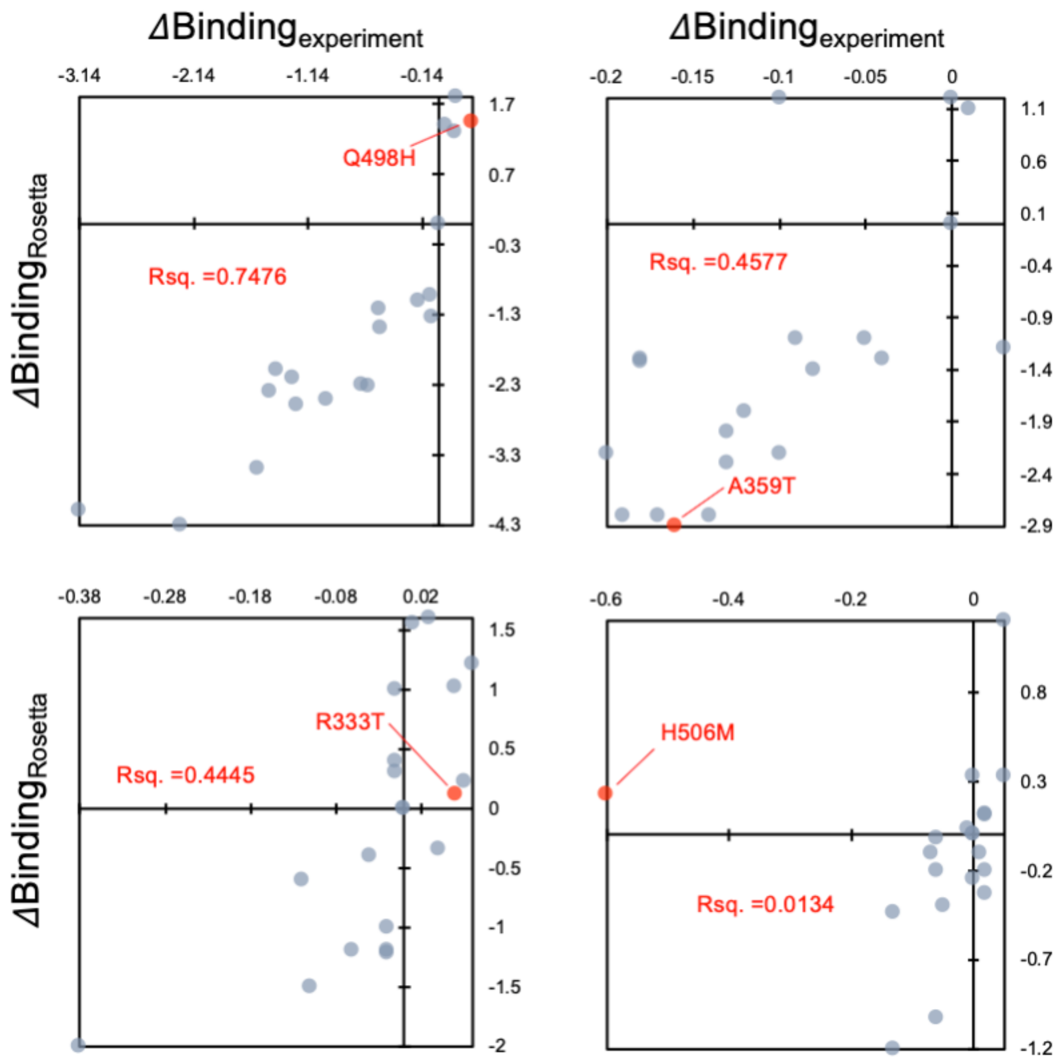


**Supplementary Figure 6. Analysis of ten independent Rosetta trajectories per mutant to report the median binding scores with hACE2 after imposing all nineteen amino acid point mutations at these four loci on the viral RBD structure.** Each grey dot represents mutation to one of the 19 other amino acids for each position. The Y and X axes show change (*Δ*) in hACE2 binding score from the default amino acid type at the backbone loci. The dot at origin (0,0), thus, represents the unmutated state (i.e. Q484H etc.). The red dot represents the mutations we reported in the previous study. We see that the R-squared correlation values are better in ¾ cases – except H505M. Each dot represents the binding score averaged over 10 independent simulation trajectories performed using Rosetta. Each simulation trajectory involves (a) making the mutation, (b) minimizing the complex between mutated RBD and hACE2, and (c) interaction energy calculation.
